# High side chain promiscuity of the terminal enzyme in the homologation pathway for L-phenylalanine and L-tyrosine

**DOI:** 10.64898/2026.06.15.732371

**Authors:** Rebecca M. Lang Harman, H. Grace Blacksone, Juan-Paolo Reynes, Anna Parviainen, Daniela Figueredo, Jorge Nochebuena, Shogo Mori

## Abstract

Natural product (NPs) and their derivatives are a major source of small-molecule drugs, and the building blocks of these NPs are often amino acids. These include both proteinogenic and nonproteinogenic amino acids (NPAAs), the latter of which expand the structural diversity of NPs. Homologation, or the addition of a methylene group to the amino acid side chain, is one modification that generates NPAAs. If the natural homologation pathway can be characterized and engineered, it could be used to diversify NPs. In this study, we investigated the terminal enzyme of this pathway, HphB, to determine its substrate scope. HphB was tested with various substrates that differed in backbone and/or side chain structures relative to its natural substrate. The results showed that HphB exhibits high promiscuity toward substrates with different side chains while maintaining strict specificity for the substrate backbone. Comparative analysis with two homologous enzymes from primary metabolic pathways revealed that HphB displays markedly higher substrate promiscuity. Bioinformatics analysis and structural modeling suggest that this promiscuity arises from the absence of a “lid” over the active site, resulting in increased solvent exposure of the substrate side chain. This study highlights the unique substrate flexibility of HphB and is a step toward engineering the homologation pathway to generate amino acid derivatives.

## INTRODUCTION

Natural products (NPs) are molecules that are synthesized by living organisms such as plants, fungi, and animals. These molecules play an integral part in drug discovery and development, as 66.7% of small molecule drugs approved between 1981 and September 2019 by the United States Food and Drug Administration (U.S. FDA) were NPs or NP-related molecules.^1^ NPs can possess many useful drug-like features that are hard to reproduce via organic chemistry methods,^2^ such as higher molecular weight, rigidity, and many stereocenters, which allow them to possess a wider range of chemical properties than typical synthetic small-molecule drugs.^3^ Identifying and characterizing the biosynthetic genes that produce these NPs or their building blocks could lead to engineering and coupling of these pathways with other methods to produce more effective NP-related drugs as well as achieving less wasteful synthesis.^2, 4^ One of the largest classes of NPs is nonribosomal peptides (NRPs), which use amino acids as their building blocks, including both proteinogenic and nonproteinogenic amino acids (NPAAs).^5^ Approximately 500 amino acids have been identified in nature, with only 20 of them being proteinogenic amino acids. This makes NPAAs critical to the diversity of NRPs.^6^ NPAAs are known to help improve drug-like properties such as half-life and bioavailability.^7^

NPAAs can be formed through modification of proteinogenic amino acids. The biosynthetic genes that are used to modify amino acids to be incorporated into NPs tend to be found within the same biosynthetic gene clusters. One example of a modification to an amino acid is homologation, which is the addition or deletion of a methylene group in the side chain; there are over 450 examples of homologated amino acids that have been found in NPs so far.^8^ Homologated amino acids are currently incorporated into several types of drugs, such as angiotensin-converting enzyme (ACE) inhibitors,^9^ but the current synthetic method of homologating some amino acids can be expensive and generate harmful waste. The natural homologation pathway for L-Phe and L-Tyr was identified in the cyanobacterium *Nostoc punctiforme* PCC73102 (Figure 1).^10^ If the homologation pathway could be characterized and engineered to homologate other proteinogenic amino acids, it would be a valuable resource for NP drug creation and modification.

**Figure 1.**
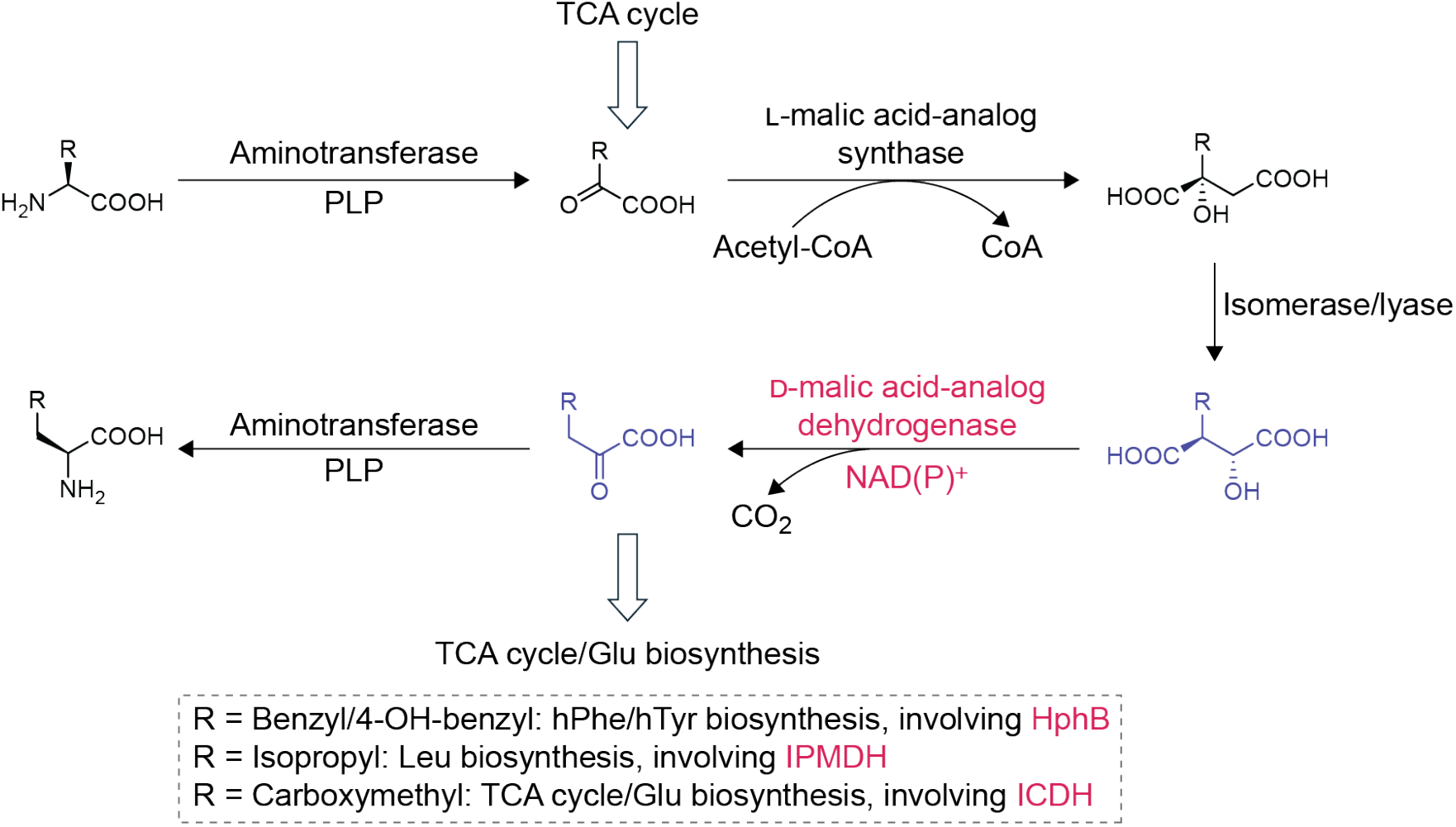
The homologation pathway of amino acids. The red-colored enzyme step is the focus of this study. For the homologation of L-Phe and L-Tyr, aminotransferase = ArAT, L-malic acid-analog synthase = HphA, isomerase/lyase = HphCD, and D-malic acid-analog dehydrogenase = HphB. Each abbreviation depicts the following: PLP = pyridoxal 5’-phosphate, CoA = coenzyme A, NAD(P)^+^ = Nicotinamide adenine dinucleotide (phosphate), TCA cycle = tricarboxylic acid cycle, IPMDH = 3-isopropylmalate dehydrogenase, ICDH = isocitrate dehydrogenase.

The homologation pathway consists of three enzymes: HphA, HphCD, and HphB, in addition to well-characterized aromatic aminotransferase (ArAT).^10^ HphA, the first enzyme in the homologation pathway, has been recently characterized as substrate-selective,^11^ and it was then demonstrated to be amenable to engineering by expanding its substrate scope.^12^ In this project, we characterized HphB, the terminal enzyme in the homologation pathway, focusing on its substrate selectivity. Because the homologation pathway is hypothesized to be homologous to the L-Leu and L-Glu biosynthesis pathways, we compared the enzymatic activity of HphB to the activities of the corresponding enzymes in the homologous pathways. The homologous enzyme to HphB in the L-Leu biosynthesis pathway is 3-isopropylmalate dehydrogenase (IPMDH; often termed as LeuB),^13^ while that in the L-Glu biosynthesis pathway is isocitrate dehydrogenase (ICDH), which is part of the tricarboxylic acid cycle.^14^ We compared their activities through establishing their substrate profiles and measuring their Michaelis-Menten kinetic parameters. We also utilized bioinformatics and modeling methods, including AlphaFold3 modeling and docking study, to support our observation: the high substrate side chain promiscuity of HphB.

## RESULTS AND DISCUSSION

### Functional evaluation of HphB, IPMDH, and ICDH from Nostoc punctiforme

To characterize HphB, the *hphB* gene found in *Nostoc punctiforme* PCC73102 was cloned and expressed in *Escherichia coli*. The HphB protein was expressed with a 6×His tag at either the N-terminus (HphB N-His) or C-terminus (HphB C-His). Taking advantage of the production of NAD(P)H from the reaction, we used spectrophotometry to monitor the reaction progress at 340 nm. Using HphB C-His, the reaction conditions were optimized using D-malic acid (D-MA; Gly counterpart of the natural substrate of HphB) as the substrate and NAD^+^ as the co-substrate based on previously reported conditions for characterizing the homologous enzyme IPMDH.^15, 16^ The activity levels of HphB N-His and C-His were compared by monitoring the reaction progress using the optimized reaction conditions, and the results indicated that HphB C-His displayed a similar, although slightly higher, activity level (Figure S2). Therefore, we used the C-His construct for all future experiments for HphB as well as for the other two enzymes studied herein.

To compare the substrate scope and kinetics of HphB with IPMDH and ICDH, the corresponding genes of IPMDH and ICDH in *N. punctiforme* PCC73102 were cloned and expressed in the same manner as that for HphB. To confirm their optimal co-substrate, these three enzymes were tested with NAD^+^ and NADP^+^. For these tests, due to the availability of the substrate, D-MA was used as the substrate for HphB and IPMDH, and isocitric acid (ICA; the natural substrate of ICDH) was used for ICDH. As expected, HphB and IPMDH were NAD^+^-dependent, and ICDH was the NADP^+^-dependent enzyme (Figure 2).

**Figure 2.**
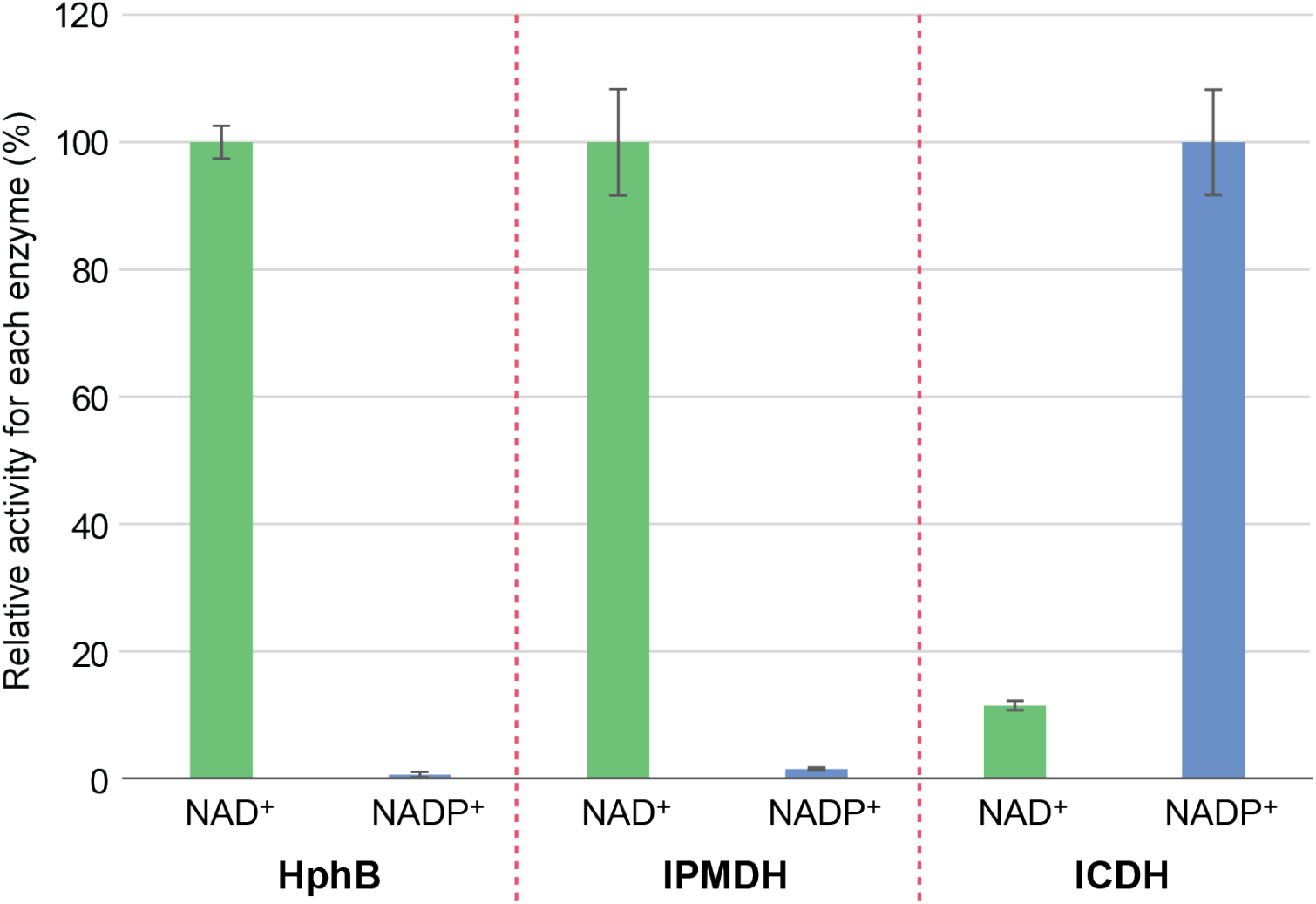
The relative activity of HphB, IPMDH, and ICDH with the co-substrate NAD^+^ and NADP^+^. The assays were duplicated. Error bars indicate the range of the data.

### Substrate profile of HphB and homologous enzymes

To determine the substrate scopes of HphB, IPMDH, and ICDH, we tested each enzyme with various substrates, including D-MA, 3-methylmalic acid (MMA), 3-isopropylmalic acid (IPMA), and ICA. We also tested HphB with L-MA, 2-hydroxybutanoic acid (HBA), and 2,4-dihydroxybutyric acid (DHBA) (Figure 3). D-MA is the Gly counterpart to the natural substrate of HphB, IPMA is the Val counterpart as well as IPMDH’s anticipated natural substrate, ICA is the Asp counterpart as well as ICDH’s anticipated natural substrate, and MMA is the Ala counterpart. L-MA has the opposite stereochemistry to D-MA at the 2-hydroxyl position. HBA and DHBA share the 2-hydroxyl acid backbone structure with other compounds, but HBA and DHBA lack the other carboxylic acid and carbonyl group of the backbone structure, respectively.

**Figure 3.**
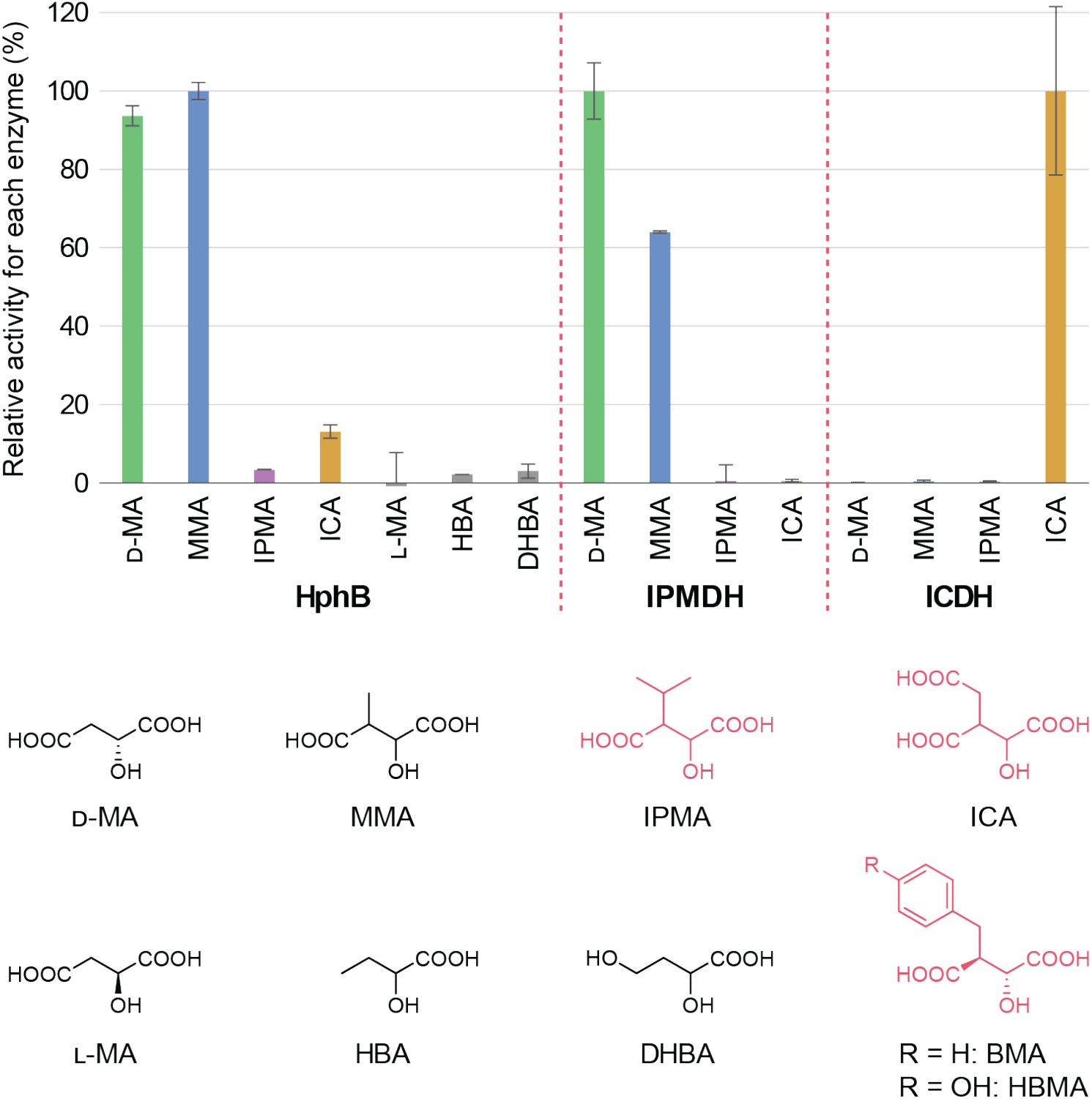
Substrate profile of HphB, IPMDH, and ICDH. The activity levels are relative to the substrate that is the greatest for each enzyme. Expected natural substrate of each enzyme is colored red. BMA/HBMA has not been tested yet. It is important to note that all tested substrates, except for MA, are racemic mixtures. Each abbreviation depicts the following: D-MA = D-malic acid, MMA = 3-methylmalic acid, IPMA = 3-isopropylmalic acid, ICA = isocitric acid, L-MA = L-malic acid, HBA = 2-hydroxybutanoic acid, DHBA = 2,4-dihydroxybutyric acid, BMA = 3-benzylmalic acid, and HBMA = 3-(4-hydroxybenzyl)malic acid. The assays were duplicated. Error bars indicate the range of the data.

The results indicate that HphB exhibits a high promiscuity toward substrates with different side chain structures than its natural substrate, 3-benzylmalic acid (BMA). However, HphB is very specific to the backbone structure; the complete D-MA backbone structure ((*R*)-2-hydroxybutanedioic acid) is required for activity. Even lack of one carbonyl group or carboxyl group almost completely disrupted the activity, as HBA and DHBA did not have significant activity. The stereochemistry at the 2-hydroxyl group is also critical, as L-malic acid did not show any activity.

Conversely, the side chain selectivity is not strict, because HphB accepted a wider range of side chains than IPMDH and ICDH. As IPMDH’s natural substrate contains an isopropyl group, the substrate with no side chain (D-MA) and an aliphatic side chain (MMA) displayed activity, while the substrate with a polar side chain (ICA) was not active at all. ICDH showed the opposite side chain selectivity to that of IPMDH; it possessed activity with the natural substrate ICA, but it did not accept any of the non-polar side chain substrates. Intriguingly, HphB demonstrated activity with both types of substrates. Although relaxed substrate selectivity of HphB was suggested by previous research,^10^ this level of promiscuity was unexpected. However, there was one critical and puzzling observation about HphB and IPMDH; the compound IPMA, thought to be the natural substrate of IPMDH, did not show activity with either HphB or IPMDH. Based on numerous previous studies on the L-Leu biosynthesis pathway and this IPMDH’s similarity to known IPMDHs, it is not reliable to conclude its inactivity. One possible explanation is that our IPMA was not a racemic mixture, and it did not include the specific stereoisomer (2*R*,3*S*) that can be recognized by the enzymes.

Overall, HphB exhibited high promiscuity to the side chain structure while maintaining strict specificity for the backbone structure. This behavior is reasonable based on its role in the homologation pathway. As the terminal enzyme, HphB operates downstream of HphA, which acts as a gatekeeper that controls the entry of substrates.^11^ Additionally, compounds with the D-MA backbone are not involved in primary metabolic pathways, and it would be advantageous for them to be converted to *α*-keto acids (e.g., pyruvic acid) that can serve as energy sources in biological systems,^17^ if they were formed. Therefore, there is limited evolutionary pressure for HphB to maintain strict selectivity toward side chain variations. In contrast, maintaining specificity toward the backbone structure is critical. For example, L-MA is one of the central intermediates involved in many primary metabolic pathways, including the tricarboxylic acid (TCA) cycle.^18^ Likewise, compounds such as HBA are derived from primary metabolites, including L-Thr.^19^ Consequently, insufficient backbone specificity could lead to the unintended processing of essential metabolites, resulting in a waste of cellular resources.

### Kinetics of HphB and homologous enzymes

We next determined Michaelis-Menten parameters for HphB with D-MA and MMA, IPMDH with D-MA and MMA, and ICDH with ICA (Table 1). These assays were performed under the same conditions used for the substrate profiling experiments, with varying substrate concentrations. Although the anticipated natural substrate of HphB has not been tested yet, the kinetic data indicate that IPMDH and ICDH exhibit substantially higher catalytic efficiency than HphB. In addition, both HphB and IPMDH showed increased efficiency with MMA relative to D-MA, consistent with the fact that their natural substrates contain larger side chains. The kinetic parameters of HphB with ICA have not been determined yet due to difficulties in obtaining consistent data. While the reaction conditions were optimized for D-MA, they may not be suitable for ICA, as aggregation of the protein was occasionally observed during these assays. This suggests that HphB with ICA is less stable under the current reaction conditions and that further optimization will be required.

**Table 1.**
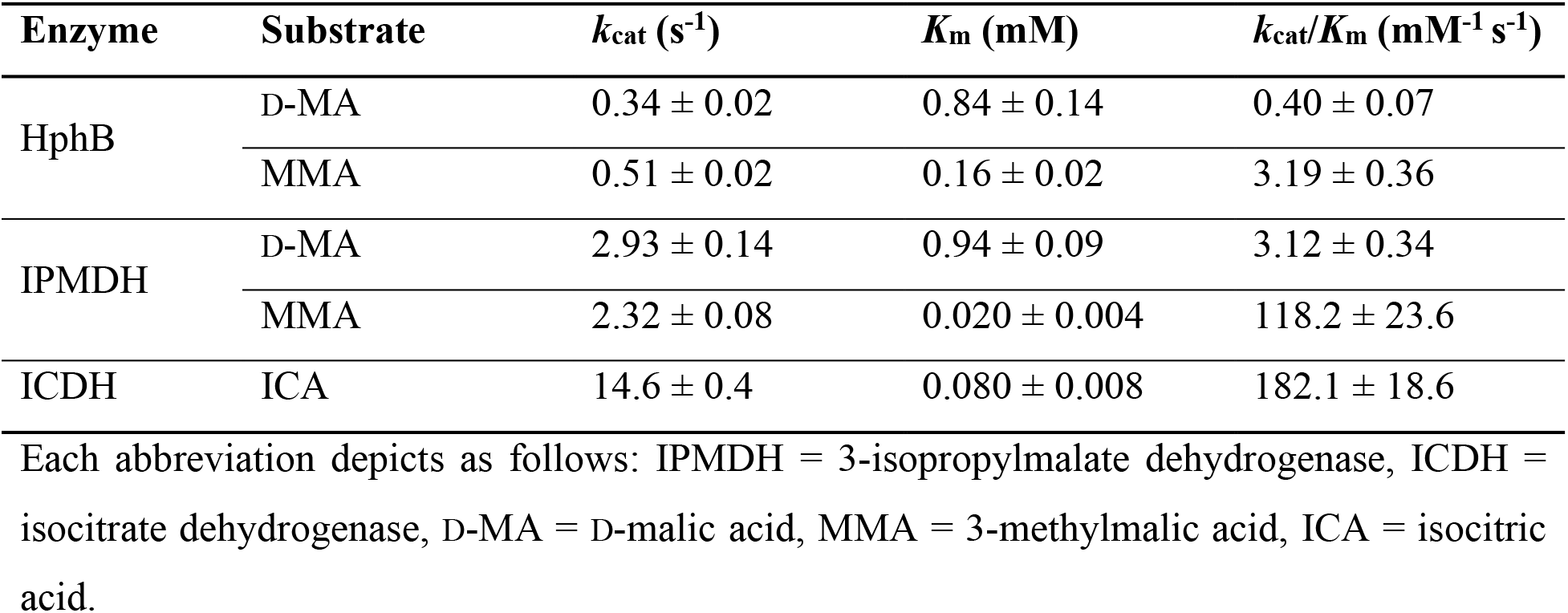
Steady-state kinetic parameters for NADH or NADPH production by HphB and homologous enzymes with the active substrate.

### Bioinformatic analysis of HphB’s active site

To identify the driver of HphB’s substrate side chain promiscuity, we conducted a bioinformatics analysis combining multiple sequence alignment and structural modeling using AlphaFold3^20^ and molecular docking.^21, 22^ First, we analyzed the amino acid sequence of HphB in comparison with IPMDHs. Although the structure of HphB has not been experimentally determined, extensive structural information is available for IPMDHs, providing a suitable reference framework. Sequence alignment with structurally characterized IPMDHs revealed a high degree of conservation between HphB and IPMDHs (Figures 4A and S4). The structures used in this analysis include those from *Shewanella benthica* DB21 MT-2 (PDB ID: 3VMK; identity/similarity = 42%/58%),^23^ *Bacillus* sp. (PDB ID: 3U1H; identity/similarity = 43%/63%),^24^ *Bacillus Coagulans* (PDB ID: 1V5B; identity/similarity = 41%/60%),^25^ *Haemophilus influenzae* Rd KW20 (PDB ID: 6XXY; identity/similarity = 41%/56%),^26^ and *Thermotoga maritima* (PDB ID: 1VLC; identity/similarity = 41%/59%).^27^ In addition to these IPMDHs, IPMDH from *N. punctiforme* PCC73102 also showed substantial identity/similarity = 37%/55%. Despite this overall conservation, one region that is distinctively different between HphB and IPMDH was identified, which is highlighted in turquoise in Figure 4A. This region consists of 14 amino acids and is highly conserved among IPMDHs, but it is absent in HphB. Analysis of one of the crystal structures bound to substrate (PDB ID: 3VMK) suggests that this region acts as a “lid” that covers the active site and stabilizes substrate binding (Figure 4B).

**Figure 4.**
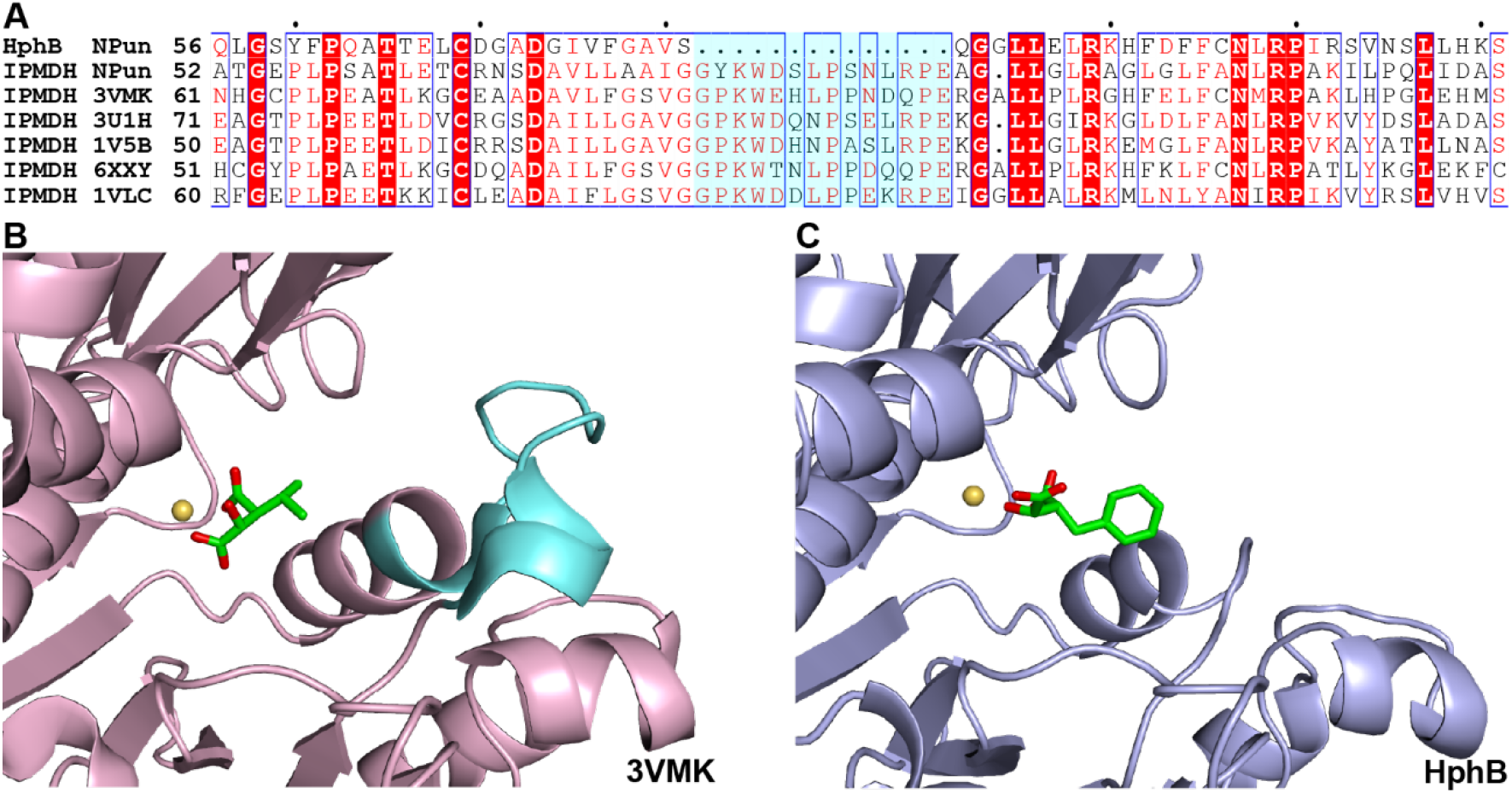
**A**. Amino acid multiple sequence alignment between HphB and IPMDH from *N. punctiforme* PCC73102 as well as structurally characterized IPMDHs, including PDB IDs: 3VMK,^23^ 3U1H,^24^ 1V5B,^25^ 6XXY,^26^ and 1VLC.^27^ **B**. The active site of the IPMDH crystal structure (PDB ID: 3VMK). **C**. AlphaFold3 model with docked BMA. The turquoise regions in Panels A and B are the amino acid sequence that lack in HphB. Yellow ball is Mg^2+^.

To further evaluate this hypothesis, we modeled the HphB structure using AlphaFold3^20^ and performed molecular docking with its proposed substrate (Figures 4C and S5). Structural comparison between the AlphaFold3 model of HphB and a homologous IPMDH structure revealed strong conservation of the overall fold and dimer architecture. Structural superposition of the HphB and IPMDH (PDB ID: 3VMK) homodimers yielded a backbone Root Mean Square Deviation (RMSD) of 1.50 Å. Consistent with previous homology-based predictions, the region proposed to function as an active-site lid in 3VMK was absent in the AlphaFold3 model of HphB, resulting in a more open and solvent-accessible substrate-binding cavity. Molecular docking of B3HB identified multiple geometrically plausible binding poses within the putative active site. The selected pose positioned the substrate hydroxyl and carboxylate groups in a chemically reasonable orientation relative to the Mg^2+^ ion while maintaining plausible interactions with surrounding active-site residues. These observations are consistent with the hypothesis that HphB possesses a more accessible substrate-binding cavity than canonical IPMDH enzymes, potentially providing a structural basis for the broader substrate scope observed experimentally.

At the same time, this open active site may reduce the strength of substrate binding, particularly for nonpolar side chains, which leads to decreased catalytic efficiency. Consistent with this interpretation, the *K*_m_ values of HphB and IPMDH with D-MA are comparable to each other (HphB: 0.84 ± 0.14 mM; IPMDH: 0.94 ± 0.09 mM), whereas the *K*_m_ value of IPMDH with MMA is approximately 10-fold lower than that of HphB (HphB: 0.16 ± 0.02 mM; IPMDH: 0.020 ± 0.004 mM). This unique structural difference likely reflects adaptation of HphB for its role in the homologation pathway of the aromatic amino acids that have larger side chains than L-Val.

## CONCLUSION

The terminal enzyme in the homologation pathway for L-Phe and L-Tyr, HphB, involved in the biosynthesis of secondary metabolites, has been characterized. It demonstrated higher side chain promiscuity than homologous enzymes in the primary metabolic pathways, including IPMDH and ICDH. Although HphB is kinetically less efficient than these homologous enzymes, it accepted substrates with both non-polar and polar side chains. The bioinformatics and computational modeling studies suggest that this promiscuity stems from lack of a “lid” over the enzyme’s active site, which is highly conserved in the IPMDH structure. Future studies should focus on testing a broader range of substrates and determining the crystal structure. These studies will guide us to engineer the homologation pathway to make it available for noncanonical amino acids, including those homologated analogs that are not readily available. The homologated amino acids would be valuable scaffolds to synthesize new drug candidates with enhanced biostability.

### ASSOCIATED CONTENT

#### Experimental Section

Full experimental methods are provided in the Supporting Information (SI). Selected methods for the key colorimetric assays are described below and are also available in Section 1.4 of the SI.

#### Colorimetric assays for HphB, IPMDH, and ICDH

The activity of HphB N-His/C-His, IPMDH C-His, and ICDH C-His enzymes was measured through colorimetric assays by taking advantage of the production of NAD(P)H. The reaction conditions were optimized using HphB C-His with D-malic acid (D-MA) from previously published conditions that were used for IPMDH.^15, 16^ The optimized reaction (100 µL) included 50 mM Tris-HCl (pH 7.5), 1 mM MnCl_2_, 100 mM KCl, 1 mM MgCl_2_, 5 mM NAD^+^ (for HphB and IPMDH) or 1 mM NADP^+^ (for ICDH). Reactions were performed on a 96-well plate at room temperature (20.6 °C), and the reaction progress was measured by a plate reader (xMark Microplate Absorbance Spectrophotometer, Bio-Rad, Hercules, CA). For kinetic assays, the concentration of produced NAD(P)H was calculated based on the standard curve for each compound (Figure S1). The standard curve was generated by measuring the absorbance at 340 nm with 0, 0.05, 0.1, 0.2, 0.3, 0.5, and 1 mM of NAD(P)H in the same buffer used for all other assays. These solutions were prepared in duplicate. The slope (*m* = ε × *l*; where ε is the extinction coefficient, and *l* is the path length) for absorbance (*A*) vs. concentration (*c*) was 1.751 mM^-1^ and 1.533 mM^-1^ for NADH and NADPH, respectively. The concentrations of NAD(P)H generated by the reaction were calculated by the following equation:

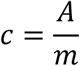

### Data Availability Statement

The data that support the findings of this study are available in the supporting information (SI) of this article in PDF format. SI Figures include the following: Standard curve for NADH and NADPH (Figure S1); Time course assays for HphB C-His *vs*. N-His (Figure S2); Kinetic assays for HphB, IPMDH, and ICDH with active substrates (Figure S3); Amino acid sequence alignment between HphB and IPMDHs (Figure S4); Superimposed structure of HphB AlphaFold3 model *vs*. IMPDH (Figure S5); and PCR primers used in this study (Table S1).

## AUTHOR INFORMATION

### Author Contributions

Study design: S.M. and R.M.L.H.; manuscript preparation: S.M. and R.M.L.H.; data generation and analysis: S.M., R.M.L.H., H.G.B., J-P.R., A.P., D.F., J.N.; Computational analysis: J.N.; funding acquisition: S.M.

## ACKNOWLEDGEMENT

The preparation of this publication was supported by the following funding sources: National Institute of General Medical Sciences of the National Institutes of Health under award number R15GM151721, start-up funds from the Department of Chemistry and Biochemistry at Augusta University, Research Scholarship and Creative Activity Program in the Division of Sponsored Programs Administration at Augusta University under award number RSCA00020, the Summer Scholars Program as well as Materials Grants from the Center for Undergraduate Research and Scholarship at Augusta University.

## References

1. Newman, D. J.; Cragg, G. M., Natural Products as Sources of New Drugs over the Nearly Four Decades from 01/1981 to 09/2019. J Nat Prod 2020, 83 (3), 770–803.

2. Butler, M. S.; La Clair, J. J., The Role of Natural Product Chemistry in Drug Discovery: Two Decades of Progress and Perspectives. J Nat Prod 2026, 89 (1), 3–28.

3. Atanasov, A. G.; Zotchev, S. B.; Dirsch, V. M.; International Natural Product Sciences, T.; Supuran, C. T., Natural products in drug discovery: advances and opportunities. Nat Rev Drug Discov 2021, 20 (3), 200–216.

4. Seshadri, K.; Abad, A. N. D.; Nagasawa, K. K.; Yost, K. M.; Johnson, C. W.; Dror, M. J.; Tang, Y., Synthetic Biology in Natural Product Biosynthesis. Chem Rev 2025, 125 (7), 3814–3931.

5. Hedges, J. B.; Ryan, K. S., Biosynthetic Pathways to Nonproteinogenic alpha-Amino Acids. Chem Rev 2020, 120 (6), 3161–3209.

6. Walsh, C. T.; O’Brien, R. V.; Khosla, C., Nonproteinogenic amino acid building blocks for nonribosomal peptide and hybrid polyketide scaffolds. Angew Chem Int Ed Engl 2013, 52 (28), 7098–124.

7. Ding, Y.; Ting, J. P.; Liu, J.; Al-Azzam, S.; Pandya, P.; Afshar, S., Impact of nonproteinogenic amino acids in the discovery and development of peptide therapeutics. Amino Acids 2020, 52 (9), 1207–1226.

8. Owens, S. L.; Ahmed, S. R.; Lang Harman, R. M.; Stewart, L. E.; Mori, S., Natural Products That Contain Higher Homologated Amino Acids. Chembiochem 2024, 25 (9), e202300822.

9. Zheng, W.; Tian, E.; Liu, Z.; Zhou, C.; Yang, P.; Tian, K.; Liao, W.; Li, J.; Ren, C., Small molecule angiotensin converting enzyme inhibitors: A medicinal chemistry perspective. Front Pharmacol 2022, 13, 968104.

10. Koketsu, K.; Mitsuhashi, S.; Tabata, K., Identification of homophenylalanine biosynthetic genes from the cyanobacterium Nostoc punctiforme PCC73102 and application to its microbial production by Escherichia coli. Appl Environ Microbiol 2013, 79 (7), 2201–8.

11. Stewart, L. E.; Owens, S. L.; Ahmed, S. R.; Lang Harman, R. M.; Mori, S., Characterization of HphA: The First Enzyme in the Homologation Pathway of l-Phenylalanine and l-Tyrosine. Chembiochem 2024, 25 (16), e202400369.

12. Lang Harman, R. M.; Blackstone, H. G.; Aruna, F. O.; Patel, S. R.; Shin, M.; NeSmith, R. K.; Dickson, D. B.; Spencer, A. C.; Mori, S., Amenability of the Gatekeeper Enzyme HphA to Engineering in the Homologation Pathway of l-Phenylalanine and l-Tyrosine through Homology-Based Site-Directed Mutagenesis. ACS Omega 2026, 11 (8), 13789–13798.

13. Kawaguchi, H.; Inagaki, K.; Kuwata, Y.; Tanaka, H.; Tano, T., 3-Isopropylmalate dehydrogenase from chemolithoautotroph Thiobacillus ferrooxidans: DNA sequence, enzyme purification, and characterization. J Biochem 1993, 114 (3), 370–7.

14. Reitzer, L., Biosynthesis of Glutamate, Aspartate, Asparagine, L-Alanine, and D-Alanine. EcoSal Plus 2004, 1 (1).

15. Yamada, T.; Akutsu, N.; Miyazaki, K.; Kakinuma, K.; Yoshida, M.; Oshima, T., Purification, catalytic properties, and thermal stability of threo-Ds-3-isopropylmalate dehydrogenase coded by leuB gene from an extreme thermophile, Thermus thermophilus strain HB8. J Biochem 1990, 108 (3), 449–56.

16. Balashova, N. V.; Zavileyskiy, L. G.; Artiukhov, A. V.; Shaposhnikov, L. A.; Sidorova, O. P.; Tishkov, V. I.; Tramonti, A.; Pometun, A. A.; Bunik, V. I., Efficient Assay and Marker Significance of NAD(+) in Human Blood. Front Med (Lausanne) 2022, 9, 886485.

17. Song, Y.; Li, J.; Shin, H. D.; Liu, L.; Du, G.; Chen, J., Biotechnological production of alpha-keto acids: Current status and perspectives. Bioresour Technol 2016, 219, 716–724.

18. Lu, J.; Zhang, S.; Wu, S.; Gao, C., l-malic acid: A multifunctional metabolite at the crossroads of redox signaling, microbial symbiosis, and therapeutic innovation. Arch Biochem Biophys 2025, 772, 110554.

19. Yu, X.; Li, Y.; Wang, X., Molecular evolution of threonine dehydratase in bacteria. PLoS One 2013, 8 (12), e80750.

20. Abramson, J.; Adler, J.; Dunger, J.; Evans, R.; Green, T.; Pritzel, A.; Ronneberger, O.; Willmore, L.; Ballard, A. J.; Bambrick, J.; Bodenstein, S. W.; Evans, D. A.; Hung, C. C.; O’Neill, M.; Reiman, D.; Tunyasuvunakool, K.; Wu, Z.; Zemgulyte, A.; Arvaniti, E.; Beattie, C.; Bertolli, O.; Bridgland, A.; Cherepanov, A.; Congreve, M.; Cowen-Rivers, A. I.; Cowie, A.; Figurnov, M.; Fuchs, F. B.; Gladman, H.; Jain, R.; Khan, Y. A.; Low, C. M. R.; Perlin, K.; Potapenko, A.; Savy, P.; Singh, S.; Stecula, A.; Thillaisundaram, A.; Tong, C.; Yakneen, S.; Zhong, E. D.; Zielinski, M.; Zidek, A.; Bapst, V.; Kohli, P.; Jaderberg, M.; Hassabis, D.; Jumper, J. M., Accurate structure prediction of biomolecular interactions with AlphaFold 3. Nature 2024, 630 (8016), 493–500.

21. Eberhardt, J.; Santos-Martins, D.; Tillack, A. F.; Forli, S., AutoDock Vina 1.2.0: New Docking Methods, Expanded Force Field, and Python Bindings. J Chem Inf Model 2021, 61 (8), 3891–3898.

22. Trott, O.; Olson, A. J., AutoDock Vina: improving the speed and accuracy of docking with a new scoring function, efficient optimization, and multithreading. J Comput Chem 2010, 31 (2), 455–61.

23. Nagae, T.; Kato, C.; Watanabe, N., Structural analysis of 3-isopropylmalate dehydrogenase from the obligate piezophile Shewanella benthica DB21MT-2 and the nonpiezophile Shewanella oneidensis MR-1. Acta Crystallogr Sect F Struct Biol Cryst Commun 2012, 68 (Pt 3), 265–8.

24. Hobbs, J. K.; Shepherd, C.; Saul, D. J.; Demetras, N. J.; Haaning, S.; Monk, C. R.; Daniel, R. M.; Arcus, V. L., On the origin and evolution of thermophily: reconstruction of functional precambrian enzymes from ancestors of Bacillus. Mol Biol Evol 2012, 29 (2), 825–35.

25. Fujita, K.; Tsuchiya, D.; Adachi, W.; Suzuki, K.; Tsunoda, M.; Minami, H.; Sekiguchi, T.; Kobayashi, N.; Mizui, R.; Tsuzaki, S.; Nakamura, S.; Takenaka, A., Crystal structure of a highly thermo-stabilized mutant of 3-isopropylmalate dehydrogenase from Bacillus coagulans: An evaluation of local packing density in the hydrophobic core. Protein Data Bank, 2003.

26. Miggiano, R.; Martignon, S.; Minassi, A.; Rossi, F.; Rizzi, M., Crystal structure of Haemophilus influenzae 3-isopropylmalate dehydrogenase (LeuB) in complex with the inhibitor O-isobutenyl oxalylhydroxamate. Biochem Biophys Res Commun 2020, 524 (4), 996–1002.

27. (JCSG), J. C. f. S. G., Crystal structure of 3-isopropylmalate dehydrogenase (TM0556) from Thermotoga maritima at 1.90 A resolution. Protein Data Bank, 2004.

