## Supplementary material for "High side chain promiscuity of the terminal enzyme in the homologation pathway for L-phenylalanine and L-tyrosine": Figure S1; Figure S2; Figure S3; Figure S4; Figure S5

**Outline:**


1. Materials and Methods — Page

1.1 Bacterial Strains, Plasmids, Materials, and Instrumentation 3

1.2 Cloning of the expression construct for HphB, IPMDH, and ICDH 3

1.3 Overexpression of HphB, IPMDH, and ICDH 4

1.4 Colorimetric assays for HphB, IPMDH, and ICDH 5

1.5 Time course assays for HphB N-His and C-His 5

1.6 Substrate profile establishment for HphB, IPMDH, and ICDH 5

1.7 Kinetic assays for HphB, IPMDH, and ICDH 6

1.8 Amino acid sequence alignment between HphB and IPMDH 6

1.9 Construction of the HphB model with AlphaFold 3 7

2. Figure and Tables

2.1 Figure S1. Standard curve for NADH and NADPH 8

2.2 Figure S2. Time course assays for HphB C-His *vs*. N-His 9

2.3 Figure S3. Kinetic assays for HphB, IPMDH, and ICDH with active substrates 10

2.4 Figure S4. Amino acid sequence alignment between HphB and IPMDHs 11

2.5 Figure S5 Superimposed structure of HphB AlphaFold3 model *vs*. IPMDH 12

2.6 Table S1. PCR primers used in this study 13

3. References

### 1. Materials and Methods

#### *1.1 Bacterial Strains, Plasmids, Materials, and Instrumentation*

Genomic DNA of *Nostoc punctiform* PCC73102 was purchased from American Type Culture Collection (ATCC 29133d-5, Manassas, VA). Chemically competent *Escherichia coli* DH5α and BL21(DE3) cells were purchased from Thermo Scientific (Waltham, MA) and Invitrogen (Waltham, MA), respectively. DNA oligonucleotides were purchased from Integrated DNA Technologies (Coralville, IA). The pET-28a overexpression plasmid for *E. coli* was purchased from Novagen (MilliporeSigma, Burlington, MA). DNA sequencing was performed by Eurofins Genomics USA (Louisville, KY). Small molecule chemicals and bacterial media were purchased from Fisher Scientific (Waltham, MA), MilliporeSigma, A2B Chem (San Diego, CA) and used without modifications. Enzymes and buffers for the molecular biology experiments were purchased from New England Biolabs (Ipswich, MA) and used as instructed.

#### *1.2 Cloning of the expression construct for HphB, IPMDH, and ICDH*

Cloning of genes coding HphB (NCBI accession #: WP_012409012), 3-isopropylmalate dehydrogenase (IPMDH; NCBI accession #: WP_012409328), and isocitrate dehydrogenase (ICDH; NCBI accession #: WP_012411727) was conducted through using polymerase chain reaction (PCR), with Phusion DNA polymerase (New England Biolabs) used as the polymerase for all PCRs. The other components for the PCRs were Phusion GC buffer, 0.2 mM dNTP, 20-200 pg/μL template, 0.5 μM forward and reverse primers, and ultra-pure water to a total volume of 50 μL. Primer pairs used for these PCRs are as follows: HphB N-His = #1/#2, HphB C-His = #3/#4, IPMDH C-His = #5/#6, and ICDH C-His = #7/#8 (Table S1). The PCR program was run with initial denaturation of 1 min at 98 °C; 30 cycles of 20 s at 98 °C, 30 s at 52 °C, and 2.5 min at 72 °C; and final extension for 5 min at 72 °C. Following PCR, the products were purified via agarose gel extraction (GeneJET Gel Extraction Kit, Thermo Fisher Scientific); the purified products were then used for double digestion with a pair of restriction enzymes NdeI/XhoI for those with N-terminus 6×His tag or NcoI/XhoI for those with C-terminus 6×His tag. The digestion products were purified via agarose gel extraction and then ligated into linearized pET-28a with T4 DNA ligase (New England Biolabs). The ligation products were transformed into chemical competent *E. coli* DH5α. A single transformed colony was culture overnight to miniprep the recombinant plasmid; each plasmid was verified through double digestion followed by sequencing reactions.

#### *1.3 Overexpression of HphB, IPMDH, and ICDH*

The HphB C-His/N-His, IPMDH C-His, and ICDH C-His constructs were overexpressed in *E. coli* and purified for *in vitro* characterization experiments. On the first day, the recombinant expression plasmid was transformed into *E. coli* BL21(DE3) cells. On the second day, 3 colonies were picked up and cultured in 3 mL of LB medium with 50 µg/mL kanamycin at 37 °C with shaking at 200 rpm until becoming cloudy, indicating the culture had reached the log phase. Then, the culture was inoculated into 1 L of LB medium with 50 µg/mL kanamycin and shaken under the same conditions until the optical density at 600 nm ($OD_{600}$) reached around 0.3-0.5. The culture was cooled to approximately 16 °C, 0.2 mM isopropyl 1-thio-β-D-galactopyranoside (IPTG) was added, and then the culture was incubated overnight at 16 °C with shaking at 200 rpm. On the third day, the cells were harvested through centrifugation at 4000 rpm for 10 min at 4 °C. Next, the cells were resuspended in approximately 30 mL of Ni-NTA binding buffer (25 mM Tri-HCl (pH 8.0), 400 mM NaCl, 5 mM imidazole, and 10% glycerol) with the addition of 1 mM phenylmethylsulfonyl fluoride (PMSF). Four cycles of sonication were performed on the resuspended cells; each cycle was 120 s with alternating 10 s "on" and 10 s "off" to disrupt the cells. Centrifugation at 14000 rpm for 45 min at 4 °C was used to remove the cell debris. Subsequently, the supernatant, which contained the solubilized protein, was incubated with Ni-NTA resin (Thermo Scientific) for >2 h at 4 °C with gentle mixing. The Ni-NTA resin was packed into a column, washed 10 times with 5 mL of wash buffer (25 mM Tris-HCl (pH 8.0), 400 mM NaCl, 40 mM imidazole, 10% glycerol), and the protein was eluted in three fractions. Each elution fraction used 1 mL of elution buffer (25 mM Tris-HCl (pH 8.0), 400 mM NaCl, 250 mM imidazole, and 10% glycerol). The first two elution fractions were dialyzed against three 1 L dialysis buffers (40 mM Tris-HCl (pH 8.0), 200 mM NaCl, 2 mM β-mercaptoethanol, and 10% glycerol); the elution fractions were dialyzed for at least 3 h in each buffer. On the fourth day, after the protein was dialyzed against the third dialysis buffer, the protein concentration was measured with a spectrophotometer at 280 nm. The protein was then flash frozen with liquid nitrogen and stored at -80 °C.

#### *1.4 Colorimetric assays for HphB, IPMDH, and ICDH*

The activity of HphB N-His/C-His, IPMDH C-His, and ICDH C-His enzymes was measured through colorimetric assays by taking advantage of the production of NAD(P)H. The reaction conditions were optimized using HphB C-His with D-malic acid (D-MA) from previously published conditions that were used for IPMDH.[1, 2] The optimized reaction (100 µL) included 50 mM Tris-HCl (pH 7.5), 1 mM $MnCl_2$, 100 mM KCl, 1 mM $MgCl_2$, 5 mM $NAD^+$ (for HphB and IPMDH) or 1 mM $NADP^+$ (for ICDH). Reactions were performed on a 96-well plate at room temperature (20.6 °C), and the reaction progress was measured by a plate reader (xMark Microplate Absorbance Spectrophotometer, Bio-Rad, Hercules, CA). For kinetic assays, the concentration of produced NAD(P)H was calculated based on the standard curve for each compound (Figure S1). The standard curve was generated by measuring the absorbance at 340 nm with 0, 0.05, 0.1, 0.2, 0.3, 0.5, and 1 mM of NAD(P)H in the same buffer used for all other assays. These solutions were prepared in duplicate. The slope ($m = \varepsilon \times l$; where ε is the extinction coefficient, and $l$ is the path length) for absorbance ($A$) vs. concentration ($c$) was 1.751 $mM^{-1}$ and 1.533 $mM^{-1}$ for NADH and NADPH, respectively. The concentrations of NAD(P)H generated by the reaction were calculated by the following equation:

$$c = \frac{A}{m}$$

#### *1.5 Time course assays for HphB N-His and C-His*

Time-course assays were performed with HphB N-His and C-His, to compare activity levels of HphB with a 6×His tag at N-terminus and C-terminus, respectively (Figure S2). Assays were performed in the same manner as in ***Colorimetric assays for HphB, IPMDH, and ICDH***. The enzyme concentrations were 1 µM and the substrate, D-MA, was 1 mM. Each trial was duplicated, and the average of the negative control was subtracted from each trial. Based on the result obtained from these assays, all enzymes utilized afterwards were C-His.

#### *1.6 Substrate profile establishment for HphB, IPMDH, and ICDH*

Time-course assays were run to establish the substrate profile for HphB, IPMDH, and ICDH, with conditions being the same as in the previous section (***Colorimetric assays for HphB, IPMDH, and ICDH***). Enzyme concentrations for testing were 10 µM for HphB, 1 µM for IPMDH, and 0.05 μM for ICDH. The tested co-substrates were $NAD^+$ and $NADP^+$ for all enzymes with the substrate D-MA for HphB and IPMDH or isocitric acid (ICA) for ICDH. The tested substrates include D-MA, isocitric acid (ICA), 3-methylmalic acid (MMA), and 3-isopropylmalic acid (IPMA). For HphB, additional substrates, L-malic acid (L-MA), 2-hydroxybutanoic acid (HBA) and 2,4-dihydroxybutyric acid (DHBA), were included. Except for malic acids, all compounds are racemic mixtures. The relative activity was defined as the absorbance at 20 min subtracting the negative control value at the same time. The negative controls were the solutions without the substrate. These assays were performed in duplicate.

#### *1.7 Kinetic assays for HphB, IPMDH, and ICDH*

Kinetic assays for active enzyme-substrate pairs were conducted to obtain Michaelis-Menten kinetic parameters. The enzyme-substrate pairs were HphB with D-MA and MMA, IPMDH with D-MA and MMA, and ICDH with ICA. Conditions were the same as in ***Colorimetric assays for HphB, IPMDH, and ICDH***. The enzyme concentrations utilized for these assays were 2 μM HphB, 0.2 μM IPMDH, and 0.05 μM ICDH. Substrate concentrations were varied between 0.01 and 2 mM. The negative controls for each enzyme were those without the substrate. The linear portion of each reaction progress curve was taken for calculating the produced NAD(P)H concentration. The obtained data was analyzed in GraphPad Prism (GraphPad Software, Boston, MA) to obtain the Michaelis-Menten parameters (Figure S3).

#### *1.8 Amino acid sequence alignment between HphB and IPMDH*

The amino acid sequence of HphB and the closely related homolog, IPMDHs, was compared by multiple sequence alignment. The amino acid sequences used for this analysis were HphB from *N. punctiforme* PCC73102, IPMDH from *N. punctiforme* PCC73102, and the FASTA sequences obtained from structurally characterized IPMDHs, including those from *Shewanella benthica* DB21 MT-2 (PDB ID: 3VMK),[3] *Bacillus* sp. (PDB ID: 3U1H),[4] *Bacillus Coagulans* (PDB ID: 1V5B),[5] *Haemophilus influenzae* Rd KW20 (PDB ID: 6XXY),[6] and *Thermotoga maritima* (PDB ID: 1VLC).[7] Multiple sequence alignment and figure generation were performed on the MultAlin web-page (Figure S4).[8]

#### *1.9 Construction of the HphB model with AlphaFold 3*

A homodimeric model of HphB was generated using AlphaFold3[9] from the full-length amino acid sequence of the enzyme. The predicted structure was structurally aligned to the crystal structure of the homologous 3-isopropylmalate dehydrogenase (IPMDH) from *Shewanella benthica* DB21 MT-2 (PDB ID: 3VMK),[3] which served as a reference for identification of the active site. A $Mg^{2+}$ ion was positioned at the corresponding metal-binding site identified from the structural alignment with 3VMK. A three-dimensional model of the proposed substrate (BMA; 3-benzylmalic acid) was generated from its SMILES representation and positioned within the active site using the location of the ligand (IPMA; 3-isopropylmalic acid) observed in the homologous enzyme structure. Local molecular docking calculations were subsequently performed in the active site of chain A using AutoDock Vina v1.2.7,[10, 11] with a cubic search space measuring 18 Å on each side and centered on the active site identified from the homologous IPMDH structure. Docking calculations were carried out with an exhaustiveness value of 16, and twenty candidate binding poses were generated and ranked according to the Vina scoring function. The final model (Affinity: -6.285 kcal/mol) was selected based on docking scores and the consistency of ligand placement relative to the $Mg^{2+}$ ion and conserved active-site residues identified from the structural alignment.

### 2. Figures and Tables

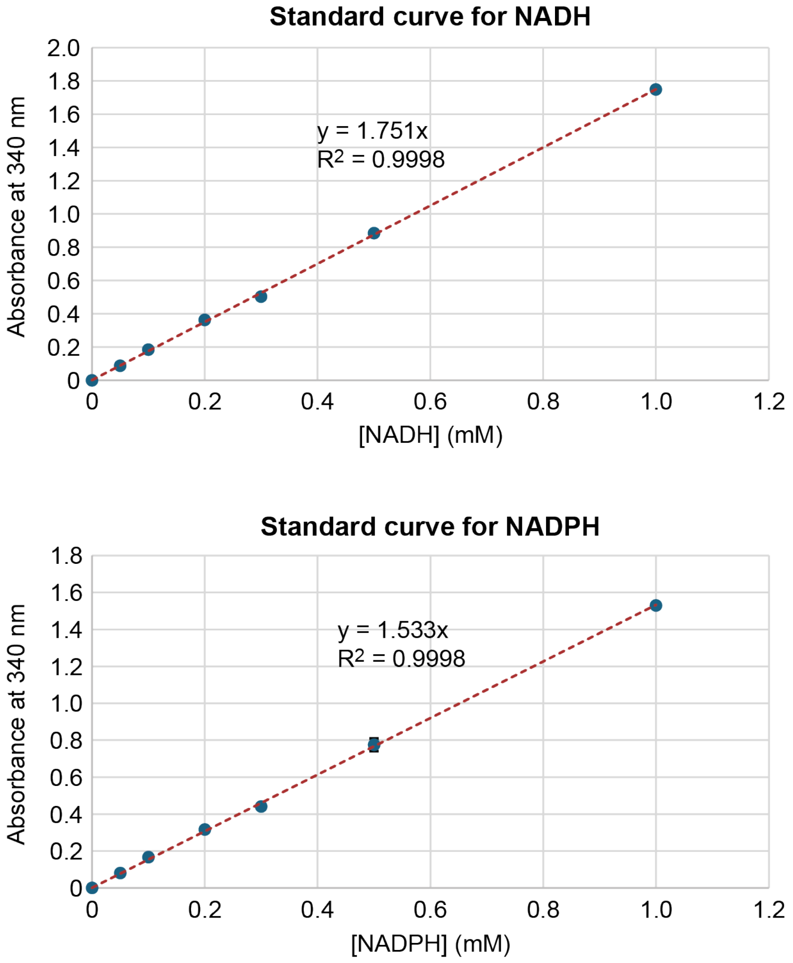


**Figure S1.** Standard curves for NADH and NADPH. The solutions were prepared in duplicate. The error bars are the range of measured values.

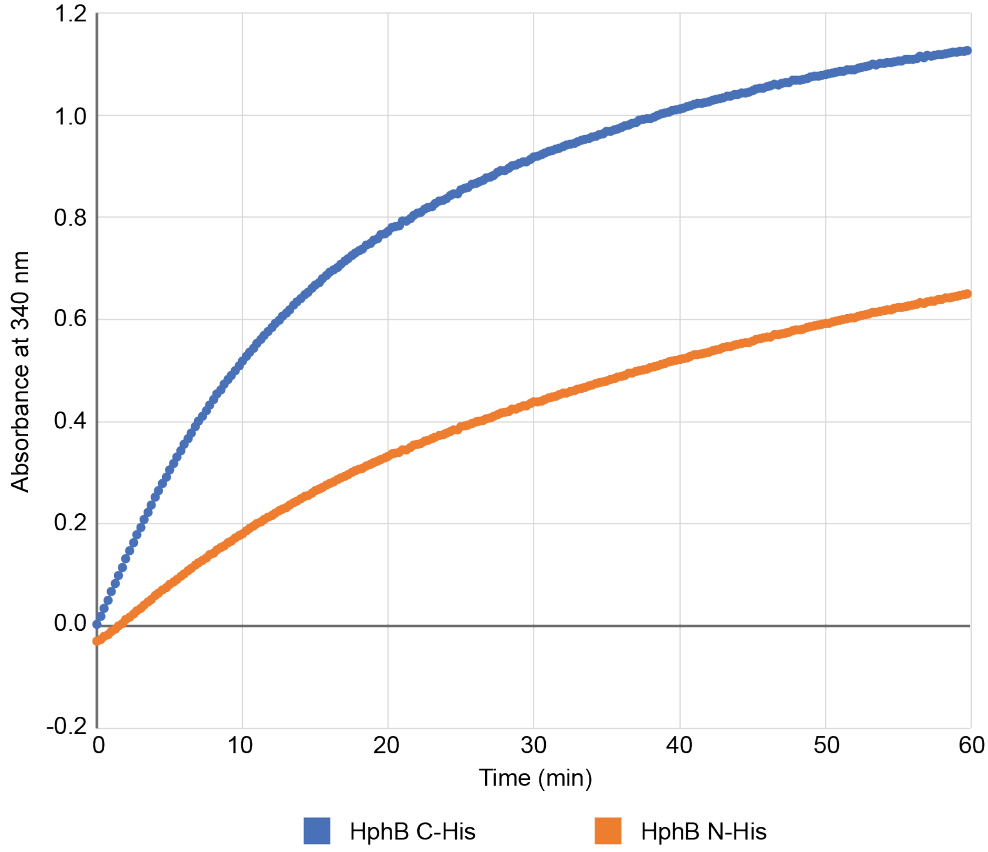


**Figure S2.** The course assays to compare HphB N-His and C-His. The assays were duplicated, and the average of each data was plotted.

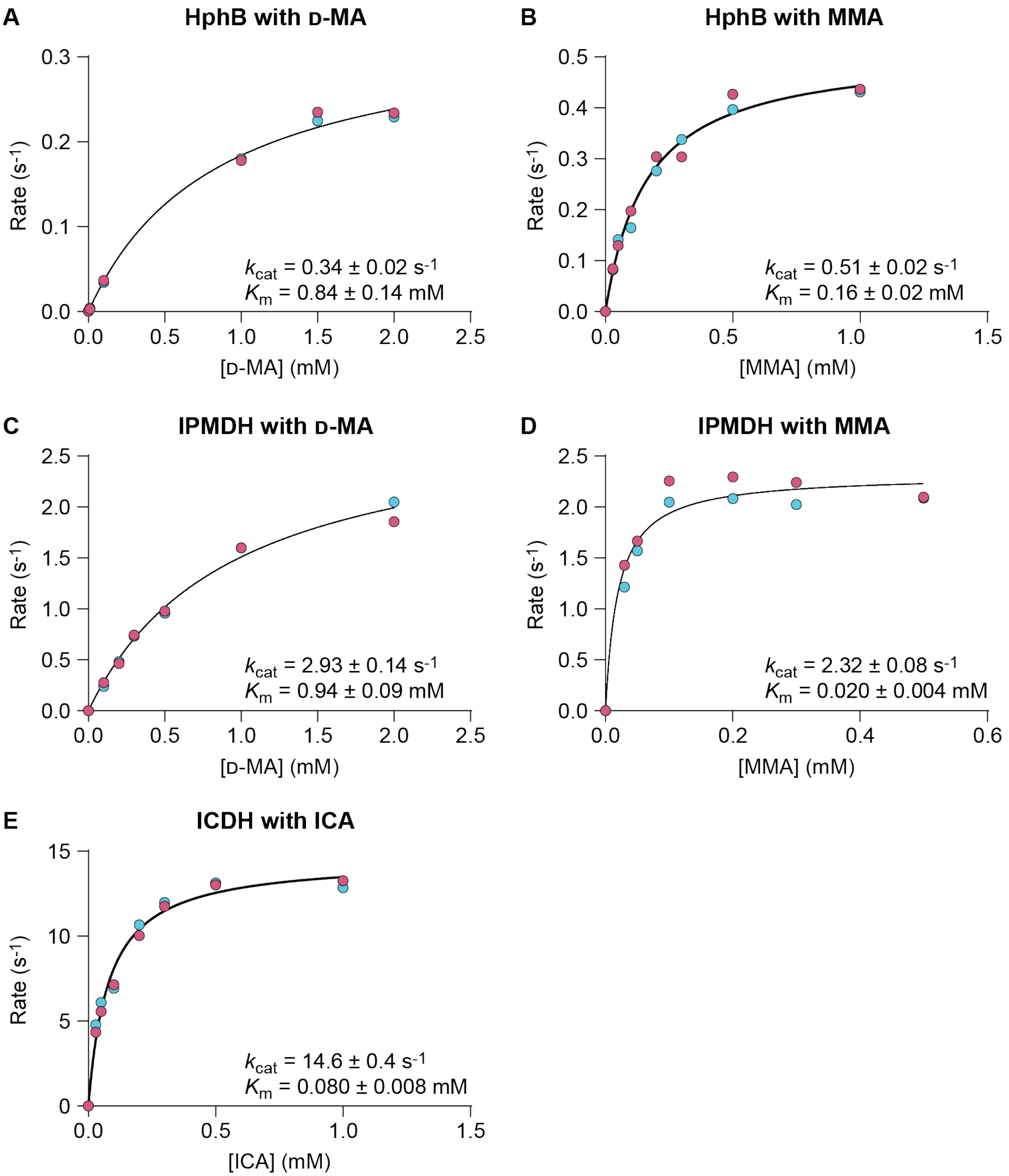


**Figure S3.** Michaelis-Menten kinetic plots for **A.** HphB with D-MA, **B.** HphB with MMA, **C.** IPMDH with D-MA, **D.** IPMDH with MMA, and **E.** ICDH with ICA. Each abbreviation depicts the following: D-MA = D-malic acid, MMA = 3-methylmalic acid, ICA = isocitric acid, IPMDH = 3-isopropylmalate dehyderogenase, and ICDH = isocitrate dehydrogenase. The assays were duplicated.

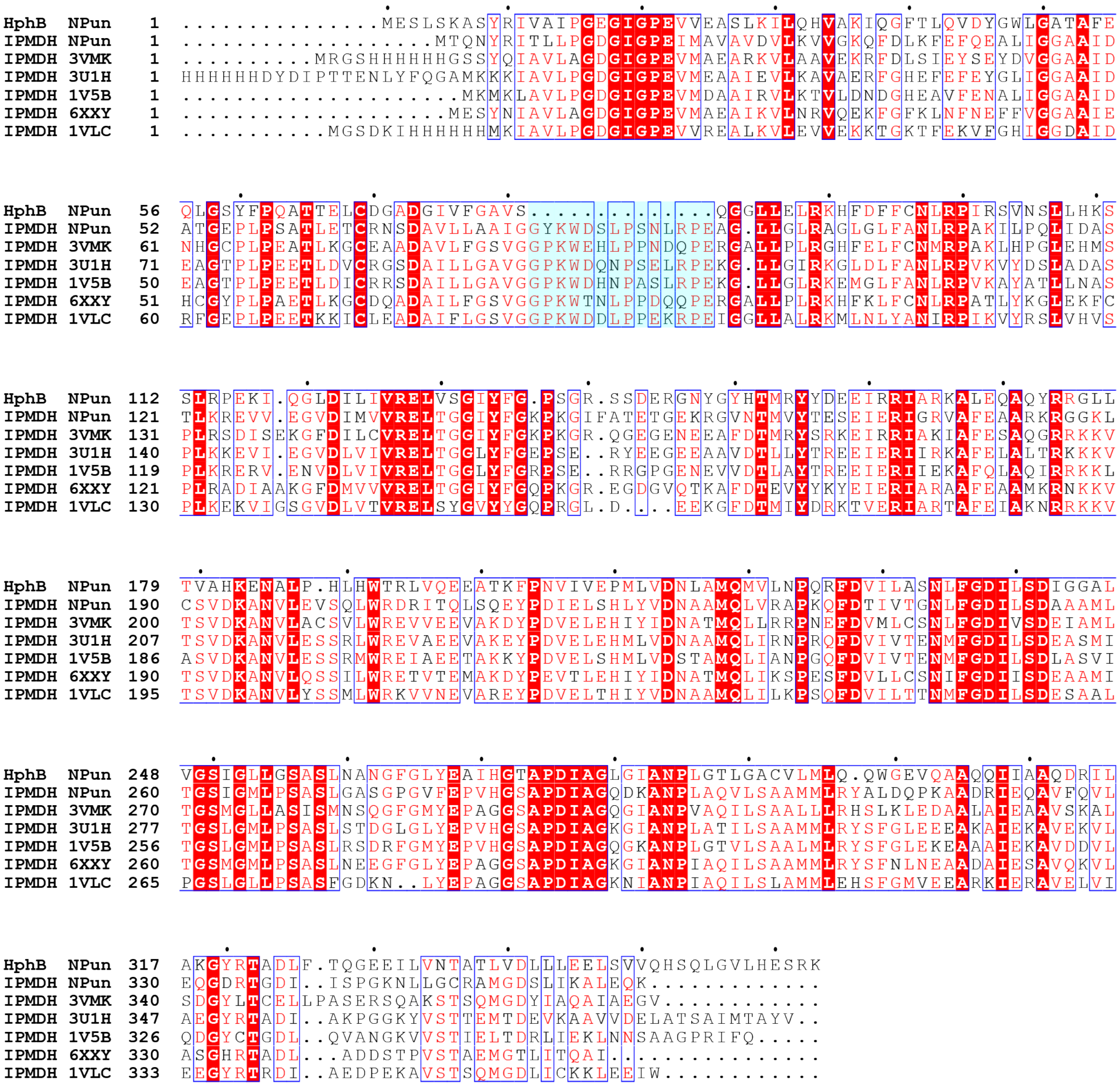

**Figure S4.** The complete amino acid sequence alignment between HphB from *N. punctiforme* PCC73102, IPMDH from *N. punctiforme* PCC73102, and structurally characterized IMPDHs. The amino acid sequence forming the “lid” of the active site that is present only in IPMDHs is shaded in turquoise.

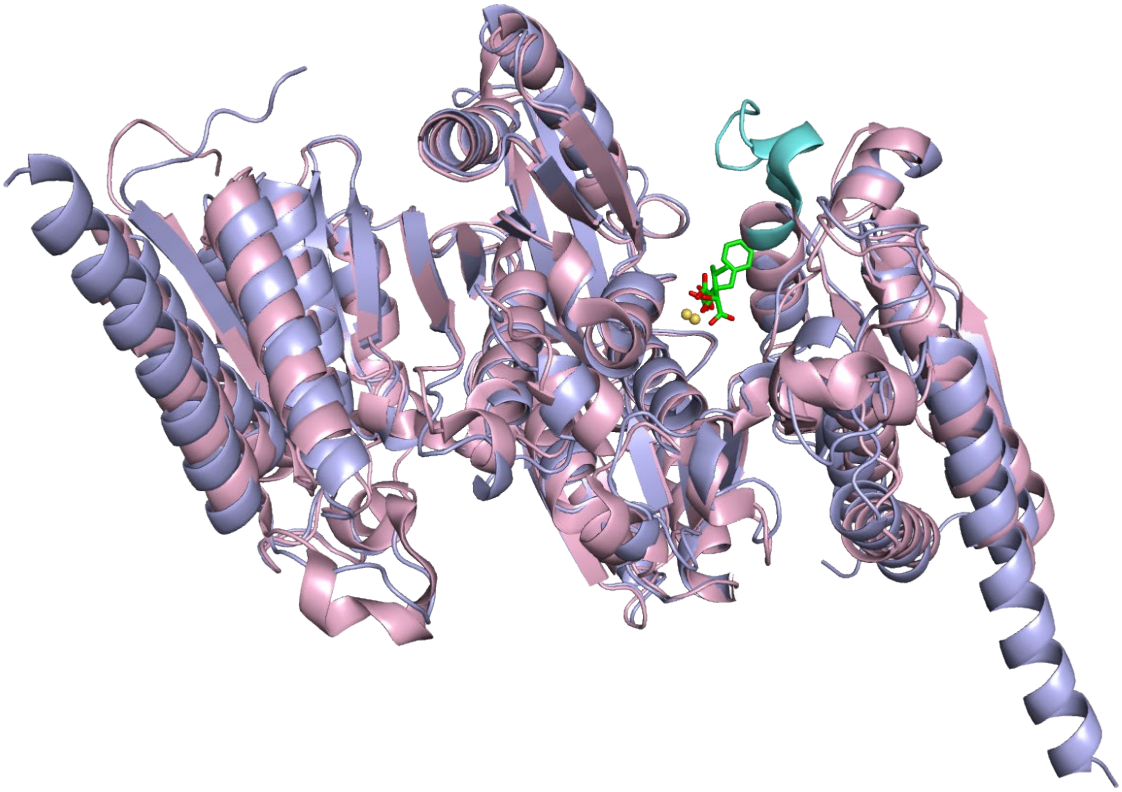

**Figure S5.** The superimposed structures of HphB and the structurally characterized IPMDH (PDB ID: 3VMK).[3] HphB is colored purple, and IPMDH is colored pink. The active site "lid" of IMPDH is colored turquoise. The substrate was shown only in the active site of the right-hand side monomer unit. The $Mg^{2+}$ is shown as a yellow ball.

**Table S1.** PCR primers used in this study

| Primer # | Primer name | Primer sequence | 5' or 3' |
|---|---|---|---|
| 1 | SM-HphB-NdeI-F | CGTGACCATATGGAATCTTTAAGTAAGGCATC | 5' |
| 2 | SM-HphB-XhoI-R-STOP | CGGTGCCTCGAGTTATTTTCGTGACTCATG | 3' |
| 3 | SM-HphB-NcoI-F | GTGACCCCATGGAATCTTTAAGTAAGGCATC | 5' |
| 4 | SM-HphB-XhoI-R-noSTOP | CGCAAACTCGAGTTTTCGTGACTCATGAAG | 3' |
| 5 | SM-NPLeuB-NcoI-F | CGTAATCCATGGGCATGACCCAGAACTACCGC | 5' |
| 6 | SM-NPLeuB-XhoI-R-noSTOP | GCCACACTCGAGTTTTTGTTCAAGAGCCTTAATG | 3' |
| 7 | SM-NPICDH-NcoI-F | GGAGTGCCATGGGCATGTACGAAAAGATTACC | 5' |
| 8 | SM-NPICDH-XhoI-R-noSTOP | CTAGTACTCGAGACCAAAATGCTGAATAATTGCG | 3' |

*Note*: The underlined nucleotide sequences are the target of the corresponding restriction enzyme.

### 3. References


1. Yamada, T.; Akutsu, N.; Miyazaki, K.; Kakinuma, K.; Yoshida, M.; Oshima, T., Purification, catalytic properties, and thermal stability of threo-Ds-3-isopropylmalate dehydrogenase coded by leuB gene from an extreme thermophile, Thermus thermophilus strain HB8. *J Biochem* **1990,** *108* (3), 449-56.
2. Balashova, N. V.; Zavileyskiy, L. G.; Artiukhov, A. V.; Shaposhnikov, L. A.; Sidorova, O. P.; Tishkov, V. I.; Tramonti, A.; Pometun, A. A.; Bunik, V. I., Efficient Assay and Marker Significance of NAD(+) in Human Blood. *Front Med (Lausanne)* **2022,** *9*, 886485.
3. Nagae, T.; Kato, C.; Watanabe, N., Structural analysis of 3-isopropylmalate dehydrogenase from the obligate piezophile Shewanella benthica DB21MT-2 and the nonpiezophile Shewanella oneidensis MR-1. *Acta Crystallogr Sect F Struct Biol Cryst Commun* **2012,** *68* (Pt 3), 265-8.
4. Hobbs, J. K.; Shepherd, C.; Saul, D. J.; Demetras, N. J.; Haaning, S.; Monk, C. R.; Daniel, R. M.; Arcus, V. L., On the origin and evolution of thermophily: reconstruction of functional precambrian enzymes from ancestors of Bacillus. *Mol Biol Evol* **2012,** *29* (2), 825-35.
5. Fujita, K.; Tsuchiya, D.; Adachi, W.; Suzuki, K.; Tsunoda, M.; Minami, H.; Sekiguchi, T.; Kobayashi, N.; Mizui, R.; Tsuzaki, S.; Nakamura, S.; Takenaka, A., Crystal structure of a highly thermo-stabilized mutant of 3-isopropylmalate dehydrogenase from Bacillus coagulans: An evaluation of local packing density in the hydrophobic core. Protein Data Bank, 2003.
6. Miggiano, R.; Martignon, S.; Minassi, A.; Rossi, F.; Rizzi, M., Crystal structure of Haemophilus influenzae 3-isopropylmalate dehydrogenase (LeuB) in complex with the inhibitor O-isobutenyl oxalylhydroxamate. *Biochem Biophys Res Commun* **2020,** *524* (4), 996-1002.
7. (JCSG), J. C. f. S. G., Crystal structure of 3-isopropylmalate dehydrogenase (TM0556) from Thermotoga maritima at 1.90 A resolution. Protein Data Bank, 2004.
8. Corpet, F., Multiple sequence alignment with hierarchical clustering. *Nucleic Acids Res* **1988,** *16* (22), 10881-90.
9. Abramson, J.; Adler, J.; Dunger, J.; Evans, R.; Green, T.; Pritzel, A.; Ronneberger, O.; Willmore, L.; Ballard, A. J.; Bambrick, J.; Bodenstein, S. W.; Evans, D. A.; Hung, C. C.; O'Neill, M.; Reiman, D.; Tunyasuvunakool, K.; Wu, Z.; Zemgulyte, A.; Arvaniti, E.; Beattie, C.; Bertolli, O.; Bridgland, A.; Cherepanov, A.; Congreve, M.; Cowen-Rivers, A. I.; Cowie, A.; Figurnov, M.; Fuchs, F. B.; Gladman, H.; Jain, R.; Khan, Y. A.; Low, C. M. R.; Perlin, K.; Potapenko, A.; Savy, P.; Singh, S.; Stecula, A.; Thillaisundaram, A.; Tong, C.; Yakneen, S.; Zhong, E. D.; Zielinski, M.; Zidek, A.; Bapst, V.; Kohli, P.; Jaderberg, M.; Hassabis, D.; Jumper, J. M., Accurate structure prediction of biomolecular interactions with AlphaFold 3. *Nature* **2024,** *630* (8016), 493-500.
10. Eberhardt, J.; Santos-Martins, D.; Tillack, A. F.; Forli, S., AutoDock Vina 1.2.0: New Docking Methods, Expanded Force Field, and Python Bindings. *J Chem Inf Model* **2021,** *61* (8), 3891-3898.
11. Trott, O.; Olson, A. J., AutoDock Vina: improving the speed and accuracy of docking with a new scoring function, efficient optimization, and multithreading. *J Comput Chem* **2010,** *31* (2), 455-61.
